# An association between *Dnmt1* and *Wnt* in the production of oocytes in the whitefly *Bemisia tabaci*

**DOI:** 10.1101/2023.09.13.557185

**Authors:** Christopher B. Cunningham, Emily A. Shelby, Elizabeth C. McKinney, Alvin M. Simmons, Allen J. Moore, Patricia J. Moore

## Abstract

The function of methylation in insects and the DNA methyltransferase (*Dnmt*) genes that influence methylation remains uncertain. We used RNAi to reduce the gene expression of *Dnmt1* within the whitefly *Bemisia tabaci*, a hemipteran species that relies on *Dnmt1* for proper gametogenesis. We then used RNA-seq to test an *a priori* hypothesis that meiosis related genetic pathways would be perturbed. We generally did not find an overall effect on meiosis related pathways. However, we found that genes in the *Wnt* pathway, genes associated with the entry into meiosis in vertebrates, were differentially expressed. Our results are consistent with *Dnmt1* knockdown influencing specific pathways and not causing general transcriptional response. This is a finding that is also seen with other insect species. We also characterized the methylome of *B. tabaci* and assessed the influence of *Dnmt1* knockdown on cytosine methylation. This species has methylome characteristics comparable to other hemipterans regarding overall level, enrichment within gene bodies, and bimodal distribution of methylated/non-methylated genes. Very little differential methylation was observed, and difference of methylation were not associated with differences of gene expression. The effect on *Wnt* presents an interesting new candidate pathway for future studies.

## Introduction

5-methylcytosine, commonly referred to as DNA methylation, is a critical epigenetic modification for many plant and animal species, including insects (Gladstad et al., 2014; Schmitz et al., 2019). DNA methylation has been linked to regulation of gene expression, transposon defense, and imprinting in vertebrates and plants (Schmitz et al., 2019). For insects, however, the role of DNA methylation is less clear. Many but not all insects possess DNA methylation and a full complement of the three DNA methylation enzyme, DNA methyltransferase 1-3 (*Dnmt1*, *Dnmt2*, *Dnmt3*; Gladstad et al., 2011; Bewick et al., 2017; Provataris et al., 2018). As with vertebrates, the genomic methylation of cytosines almost exclusively occurs within a CpG context for insects (Gladstad et al., 2016; Schmitz et al., 2019). However, insects diverge from vertebrates in that they have generally lower levels of genomic methylation and it is enriched within gene bodies and not gene promoters or transposons. Differences in cytosine methylation level are not associated with differences in gene expression within a locus across insects, another contrast to vertebrates (Moradin and Brendel, 2021; Dixon and Matz, 2022; Duncan et al., 2022; Malezka and Kucharski, 2022). There is also considerable variation among both the genomic methylation level and the gene copy number of each *Dnmt* among insect species (Gladstad et al., 2011; Bewick et al., 2017; Provataris et al., 2018; Duncan et al., 2022). Some insects have little to no detectable cytosine methylation, *Drosophila melanogaster* and *Tribolium castaneum* being prominent examples (Zemach et al., 2010; Bewick et al., 2017; Provataris et al., 2018). Others have zero annotated copies of DNA methyltransferases, as found in several species of Diptera and Strepsiptera (Gladstad et al., 2011; Bewick et al., 2017; Provataris et al., 2018; Duncan et al., 2022). There isn’t a perfect correlation between the presence of *Dnmt* genes and cytosine methylation. *Tribolium castaneum* requires *Dnmt1* function, despite not having detectible cytosine methylation in its genome (Schulz et al. 2018). Despite the general conservation of genomic cytosine methylation and its machinery, no general function of DNA methylation has been established across insects (Dixon and Matz, 2022; Duncan et al., 2022; Malezka and Kucharski, 2022).

While no general function of cytosine methylation has been established, some functions of the DNA methylation enzymes have been characterized for several species, particularly involving *DNA methyltransferase 1*. As with vertebrates *Dnmt1* is responsible for maintaining methylation patterns after a cell division (Law & Jacobsen, 2010; Gladstad et al., 2016; Schmitz et al., 2019). Reduction of *Dnmt1* expression reduces genomic cytosine methylation level following mitosis, as expected (Bewick et al., 2019; Amukamara et al., 2020; Ventos-Alfonso et al., 2020; Washington et al., 2021; Ivasyk et al. 2023). In addition to its canonical function, several studies have established that *Dnmt1* has additional functions for insects. When *Dnmt1* expression is reduced, females show reduced egg production, males show reduced sperm production, and embryos show reduced viability after oviposition for several insect species (*Bemisia tabaci*, Shelby et al., 2023; *Blattella germanica*, Ventos-Alfonso et al., 2020; *Bombyx mori*, Li et al., 2019; *Harmonia axyridis,* Gegner et al., 2020; *Oncopeltus fasciatus*, Bewick et al., 2019, Amukamara et al., 2020, Washington et al., 2021, Cunningham et al., 2023; *Ooceraea biroi*, Ivasyk et al. 2023; *Nasonia vitripennis*, Zwier et al., 2012, Arsala et al., 2022; *Nilaparvata lugens*, Zhang et al., 2015; *Tribolium castaneum*, Schulz et al., 2018); an effect also observed for mammals (*Mus musculus*, Takada et al., 2021). Somewhat unexpectedly, this effect appears to be specific to gametes as the gonads themselves do not show any gross signs of apoptosis (Amukamara et al., 2020, Washington et al., 2021; Shelby et al., 2023; Ivasyk et al. 2023). Lifespan is generally not reduced despite a reduction of *Dnmt1* expression (Amukamara et al., 2020; Shelby et al., 2023; Ivasyk et al. 2023). Gamete-specific effects are even present for *Tribolium castaneum*, a species that has functioning *Dnmt1* but no cytosine methylation (Schulz et al., 2018). These phenotypes and the apparent cell type-specific effects lead us to further investigate the genetic mechanisms that underpin the deficits, focusing on potential genetic networks.

In this study we used RNA interference (RNAi) to test the function of *Dnmt1* of the whitefly *Bemisia tabaci* (Gennadius). *Bemisia tabaci* is a global pest species (Stanley & Naranjo, 2012). As such, work has been done to understand its reproduction and the molecular pathways that underpin its reproduction with an eye towards more efficient control of this pest (Upadhyay et al., 2016; Luo et al., 2017; Grover et al., 2018; Shelby et al., 2020; Hu et al., 2022; Shelby et al., 2023). As with other insect species, reduction of *Dnmt1* expression of this species leads to greatly reduce oocyte production and embryo inviability (Shelby et al., 2023). From previous studies, gametogenesis at the mitosis-meiosis transition is arrested (Amukamara et al., 2020; Washington et al., 2021; Cunningham et al., 2023; Ivasyk et al. 2023). This led us to hypothesize that reducing *Dnmt1* expression targets specific gene networks. To that end our first aim was to test if reducing *Dnmt1* expression preferentially perturbs meiosis-related pathways, but not mitotic or general cell cycling pathways. We tested this in two ways. First, we tested a panel of *a priori* candidate genes canonically associated with each process to contrast; general cell proliferation, cell cycling, mitosis, and meiosis. Second, we tested a panel *a priori* Gene Ontology Terms representing these processes and assessed if they were enriched among the most differentially expressed genes. Finally, we assessed the general patterns of differential gene expression to explore the dataset for perturbed pathways. Our analysis showed that reducing *Dnmt1* expression does not cause a general transcriptional response but uncovered *Wnt* as a novel candidate pathway. Our second aim for this study was to characterize the general patterns of genomic cytosine methylation of this species, the effect of *Dnmt1* reduction on cytosine methylation, and the association between cytosine methylation and gene expression. We found that *B. tabaci* has cytosine methylation at a level comparable to other hemiptera. However, reducing *Dnmt1* expression did not cause pervasive reduction of cytosine methylation, but cell type-specific effects remain to be assessed for future studies. The differences of cytosine methylation that are detectable are not associated with difference of gene expression. Future experiments assessing more tissue-specific level expression is now needed to determine if and how *Dnmt1* interacts with meiotic pathways.

## Results

### *Dnmt1* RNAi knockdown

Consistent with Shelby et al. (2023), our fed RNAi treatment reduced the expression of *B. tabaci*’s single copy of *Dnmt1* within adult females (LOC109040312; log_2_ fold change: -0.0216, P = 0.0276). This is the same magnitude of knockdown that was seen with Shelby et al. (2023) when we first characterized the reproductive deficits that *Dnmt1* RNAi knockdown produced.

### *a priori* candidate gene screen

Our hypothesis that meiotic-specific candidate genes would be preferentially perturbed was not strongly supported; none of the 5 meiosis-specific candidate genes were differentially expressed (Table 1). Four of the 22 candidate genes were statistically differentially expressed after FDR thresholding (Table 1). These genes were *Wnt*-like genes, along with one *CDC* and one *SMC3*. All gene assessed both here and by Shelby et al. (2023) had the same directional change and rank order of effect sizes.

**Table 1.**
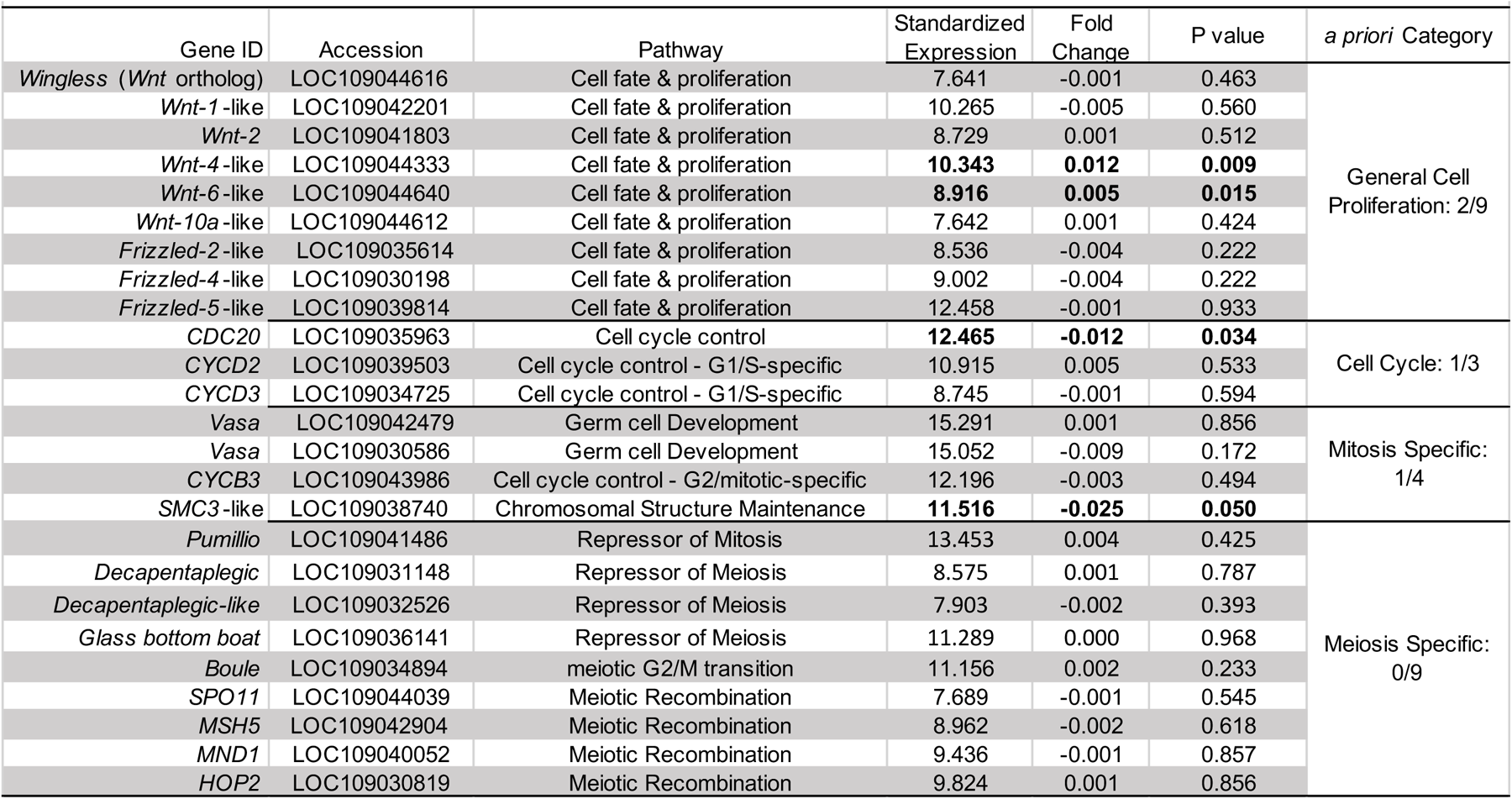
*a priori* candidate genes tested for differences between Control and *Dnmt1* knockdown treatments. Expression value and fold changes are from DESeq2. Bolded values are statisitcally significant after adaptive linear two-step procedure (Benjamini et al., 2006). Gene orthology and nomeclature is from OrthoDB.

### *a priori* GO term enrichment

Our hypothesis that the meiosis pathway would be the most affected by *Dnmt1* knockdown was supported in a minority of our *a priori* GO term enrichment analysis. In particular, the meiosis GO Term was not enriched. However, five of the six *a priori* GO terms were statistically significantly. These were among the genes that showed the least differences of expression between the control and RNAi-treatments, the opposite pattern predicted (Table 2).

**Table 2.** *a priori* GO term enrichment. Loss of *Dnmt1* does not preferentially target our predcited pathways. NES = normalized enrichment score.

|  |  |  |  |  |
| --- | --- | --- | --- | --- |
| Pathway | GO Term | Number of Annotated Genes | NES | FDR-adjusted P-value |
| Cell Cycle | GO:0007049 | 384 | -4.742 | < 0.001 |
| Gametogenesis | GO:0007276 | 681 | -4.260 | < 0.001 |
| Mitotic Cell Cycle | GO:0000278 | 255 | -4.315 | < 0.001 |
| Meiotic Cell Cycle | GO:0051321 | 144 | -3.385 | < 0.001 |
| Oogenesis | GO:0048477 | 491 | -3.644 | < 0.001 |
| Apoptotic Process | GO:0006915 | 61 | 0.676 | 8.605E-01 |

### Differential gene expression

We assessed the impact of *Dnmt1* knockdown to identify genes that were differentially expressed (11 biological replicates of *Dnmt1* and 12 of *eGFP*). The individual samples of each treatment clustered together, but the treatments were differentiated using a PCA and not statistically significantly different as the 95% confidence intervals were highly overlapping (Fig. 1). There were 96 genes (49 up-regulated, 47 down-regulated) between the treatments.

**Fig 1.**
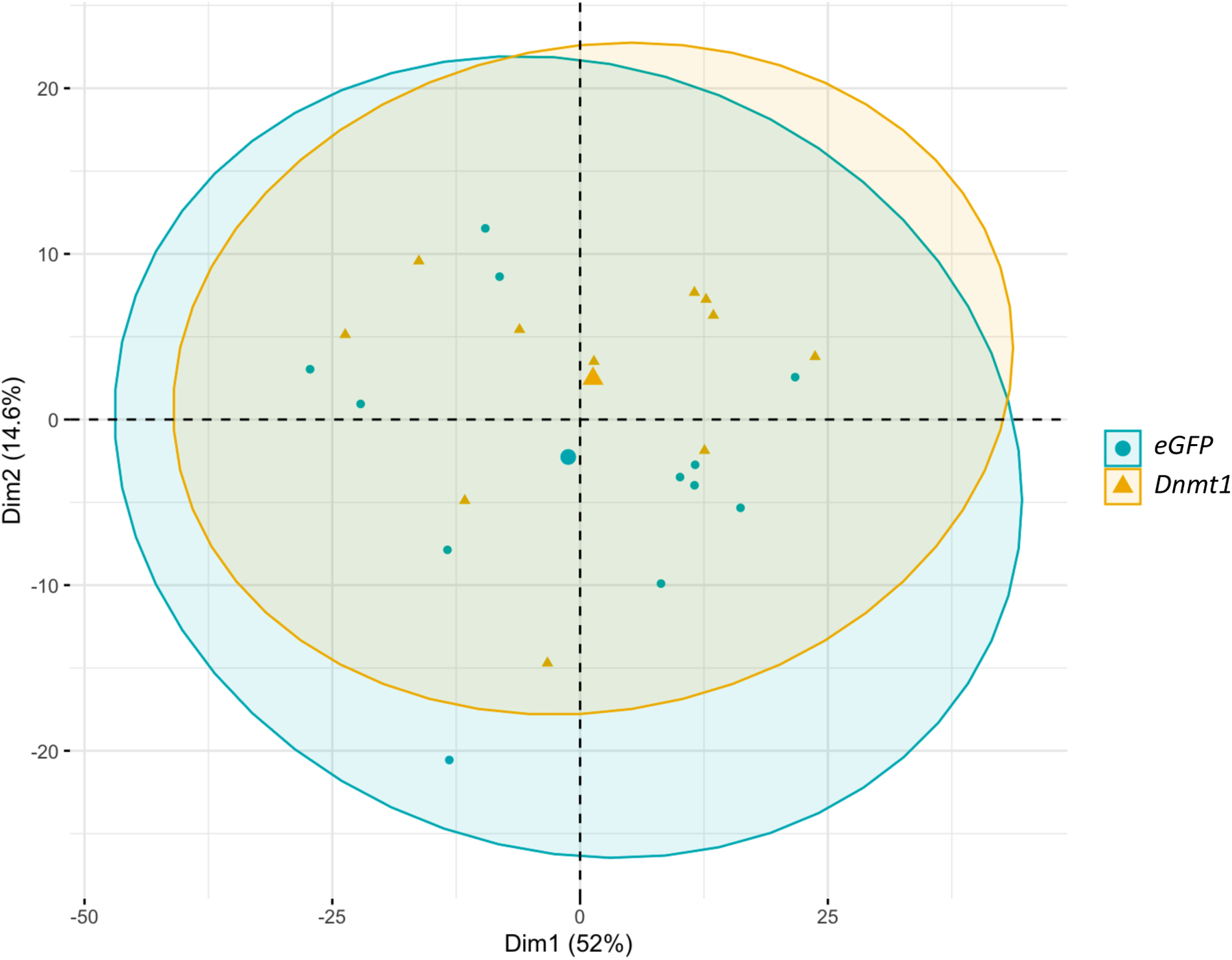
There was little differentiation of samples reflecting how few genes were differentially expressed between treatments. The percent of variation capture by each axis is in parenthesis. Smaller points represent individual samples and are colored by their treatment category (n = 12 for *eGFP*, 11 for *Dnmt1*). Large points represent the centroid (i.e., mean) of each treatment. Circles represent 95% confidence intervals.

### GO term overrepresentation

The differentially expressed genes between *Dnmt1* knockdown and controls produce 87 overrepresented GO terms. The top three Biological Process terms are female meiosis I – GO:0007144, sperm axoneme assembly – GO:0007288, and negative regulation of imaginal disc growth – GO:0045571 (P = 0.00021. P= 0.00044, P = 0.00075, respectively). Although meiosis is a top GO term, the overall list does not contain other terms related to meiosis. Specifically, the list contains only one meiosis specific term, but five terms specific to mitosis and five terms specific to general cell development (Supplementary File 3). The top three Molecular Function terms with more than one annotated gene are mRNA 3’-UTR binding – GO:000373, ribonucleoside-diphosphate reductase activity – GO:000474, and coenzyme A transmembrane transporter activity – GO:0015228 (all P = 0.0101). The top three Cellular Compartments terms with more than one annotated gene are MKS complex – GO:0036038, mitotic spindle pole – GO:0097431, and mitochondrial respiratory chain complex IV – GO:0005751 (P = 0.0108, P = 0.0162, P = 0.0162, respectively). The remainder of the terms can be found in Supplementary File 3.

### Characterization of cytosine methylation

We assessed genomic distribution and abundance of cytosine methylation to try to understand its potential function for *B. tabaci*. Cytosine methylation is found almost exclusively within a CpG context, which is consistent with other hempiterans and insects in general. Cytosines are methylated symmetrically (i.e., both strands of a CpG are methylated or not; Fig. 2), which is an indication that *Dnmt1* functions as expected. GC content of genes is higher than the background intragenic genome. This is expected because *B. tabaci* is an AT biased genome overall (genome: 39% GC, genes: 44% GC; Fig. 3A). Methylated cytosines within genic regions mirror the GC content of genes (Fig. 3B). Genes have a very bimodal distribution of methylation status (i.e., genes are either highly methylated or not; Fig. 3C). Transposons did not have consistent or high levels of methylation (Fig. 3D). Methylated cytosines are enriched within genic and exonic regions but are seen at genomic levels within introns (Fig. 4).

**Fig 2.**
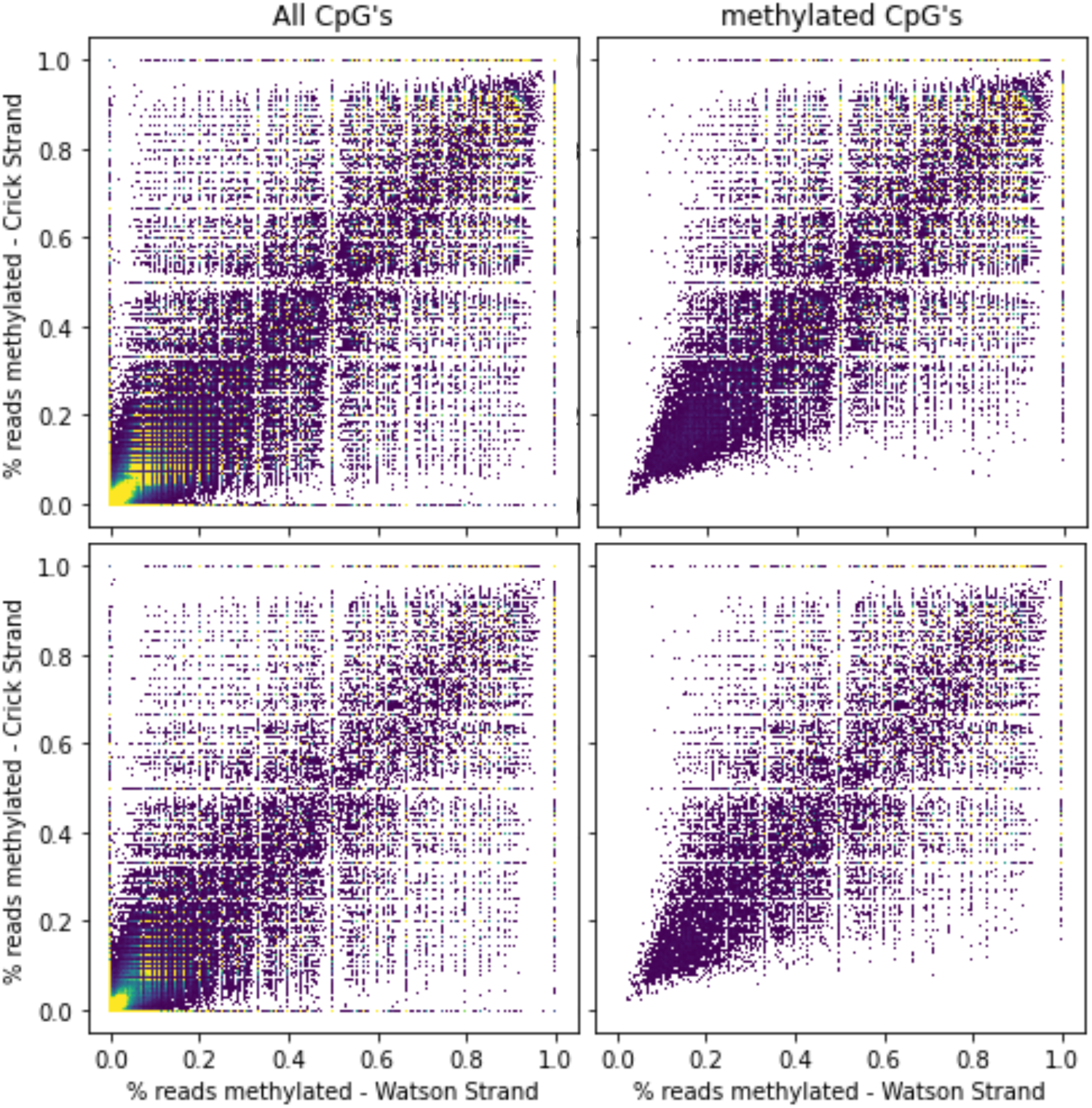
Cytosine methylation of *Bemisia tabaci* is highly symmetric, consistent with many other insects. Density plot of CpG’s methylation within different contexts. Lighter colors represent denser value regions.

**Fig 3.**
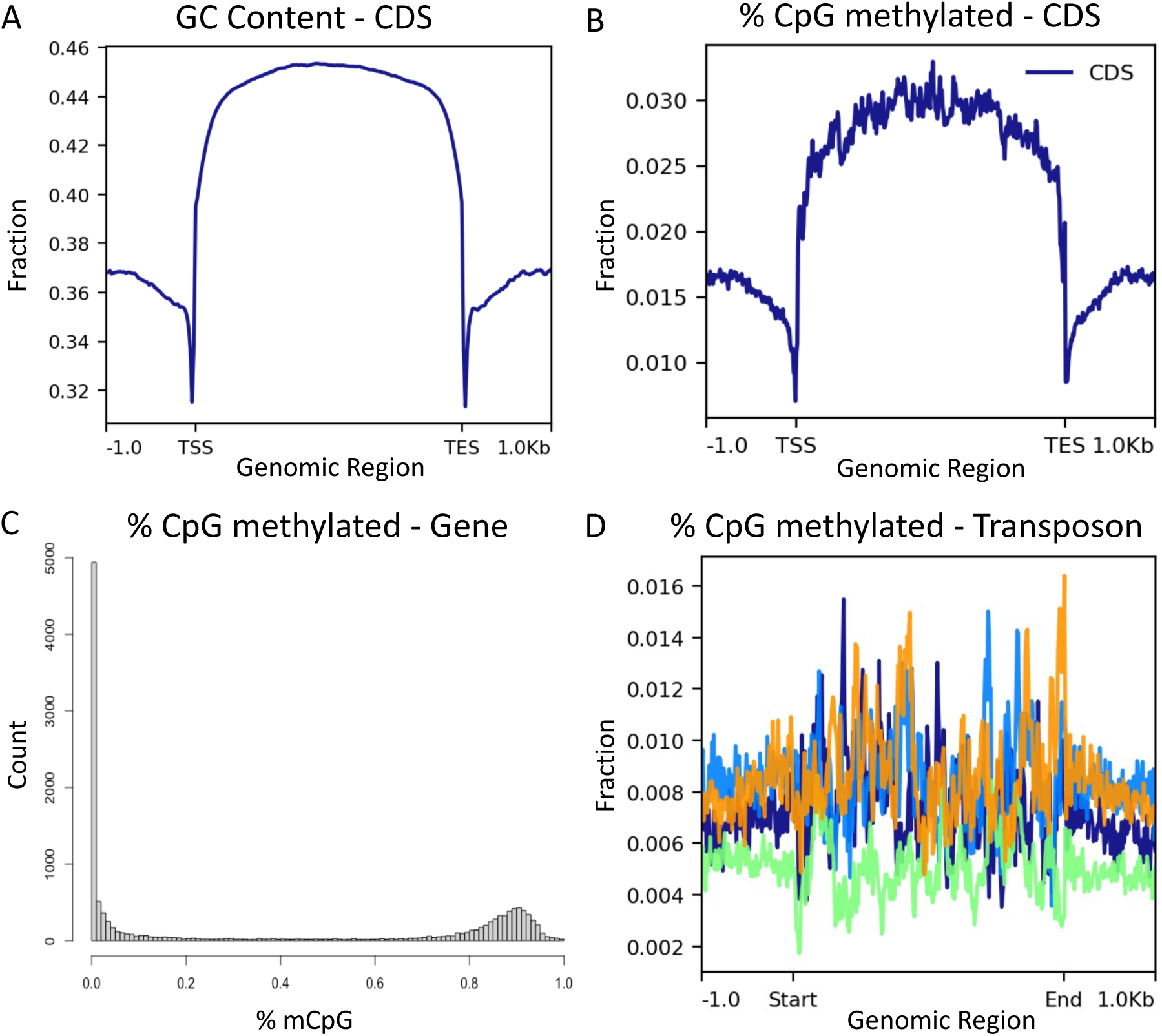
Cytosine methylation of *Bemisia tabaci* is mostly consistent with many observed trends of other insects. (A) The GC content increases over gene bodies. CDS = coding sequence, TSS = transcriptional start site, TES = transcriptional end site. (B) CpG methylation is mostly found in gene bodies. (C) Genes are either highly methylated or non-methylated. This is a more bimodal distribution than is seen for other insects. (D) There is little cytosine methylation over the most frequent transposon within the *B. tabaci* genome. Each colored line represents the four most populous families of transposons within the *B. tabaci* genome (orange = BEL, green = Gypsy, blue = hAT, navy = Mariner).

**Fig 4.**
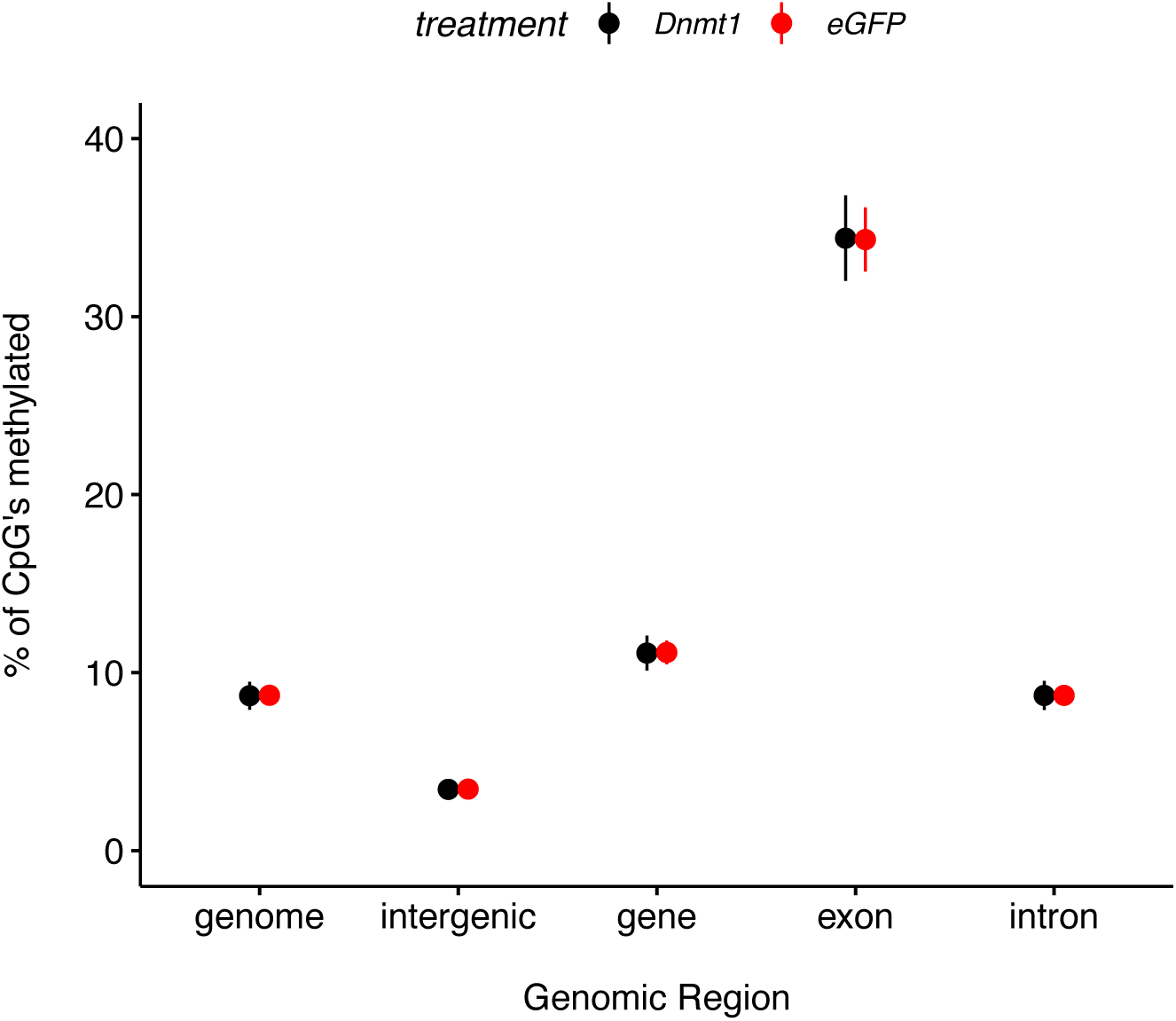
Cytosine methylation of *Bemisia tabaci* is not detectably influenced by knockdown of *Dnmt1*. Cytosine methylation is enriched with genic and exonic regions but is not detectably influenced by *Dnmt1* RNAi. Each point represents the mean ± SEM of each treatment within the labeled category.

### Cytosine methylation is associated with overall gene expression (between loci differences)

High levels of gene methylation are associated with high levels and low variances of gene expression (Fig. 5). This is a well-established pattern that suggests if cytosine methylation is performing a function, it is likely similar between *B. tabaci* and other insects.

**Fig 5.**
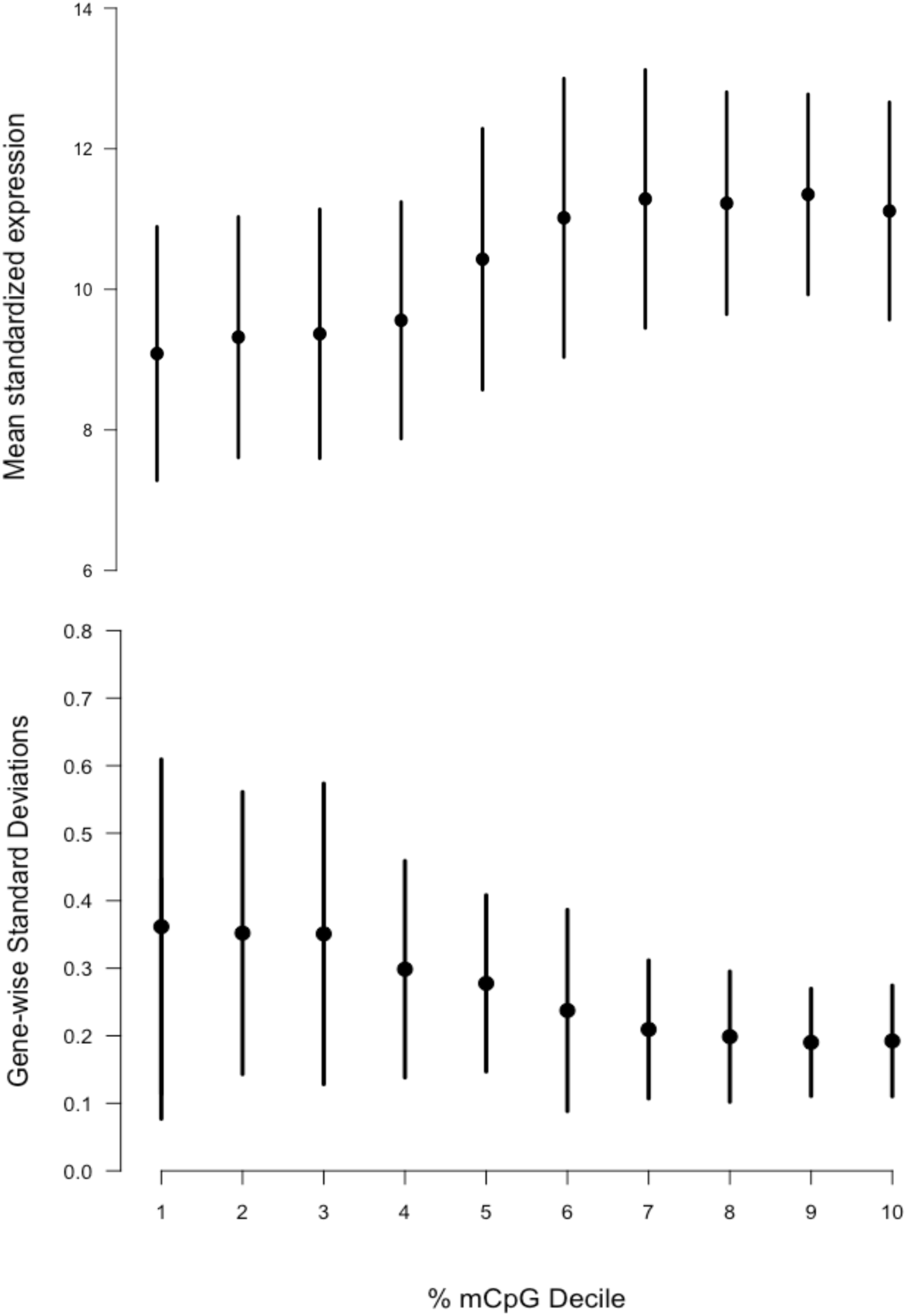
Gene body methylation is weakly, positively associated with overall gene expression levels and negatively associated with gene expression variance. (A) Plot of mean standardized gene expression values against decile of the percent of CpG’s methylated within a gene body. (B) Plot of gene-wise standard deviation against the decile of the percent of CpG’s methylated within a gene body. Both panels have values that represent the mean ± SEM.

### Differential gene methylation

After assessing the general trends of cytosine methylation within *B. tabaci*, we assessed the effect of *Dnmt1* knockdown on gene methylation level between *Dnmt1* and *eGFP* samples (five biological replicates of each treatment). Very few gene loci differed at all between the treatments; only 2,391 differed by 1% or more (Fig. 6). No gene was statistically significantly different after FDR correction.

**Fig 6.**
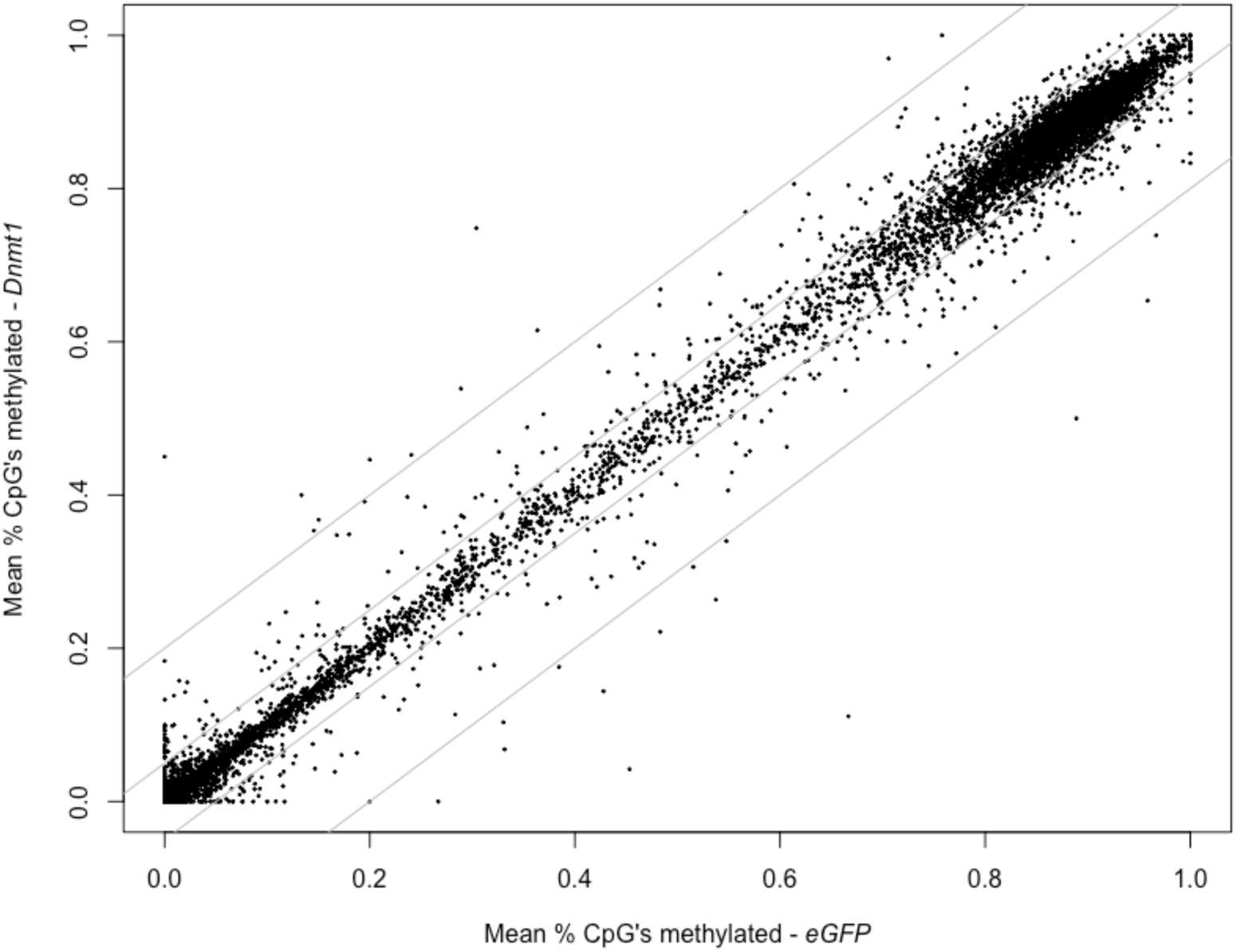
Very few genes differ for methylation status between treatments. The inner and outer diagonal lines represent 5% and 20% differentiation of *Dnmt1* vs *eGFP* treatments, respectively. Each circle represents an individual gene.

### Differential gene methylation is not associated with differential gene expression (within locus differences)

Lastly, we assessed if the difference of gene methylation that were present influenced gene expression within (and not between as above) a gene. There was no indication that difference of methylation influenced difference of gene expression (Fig. 7). This is consistent with almost all other insects.

**Fig 7.**
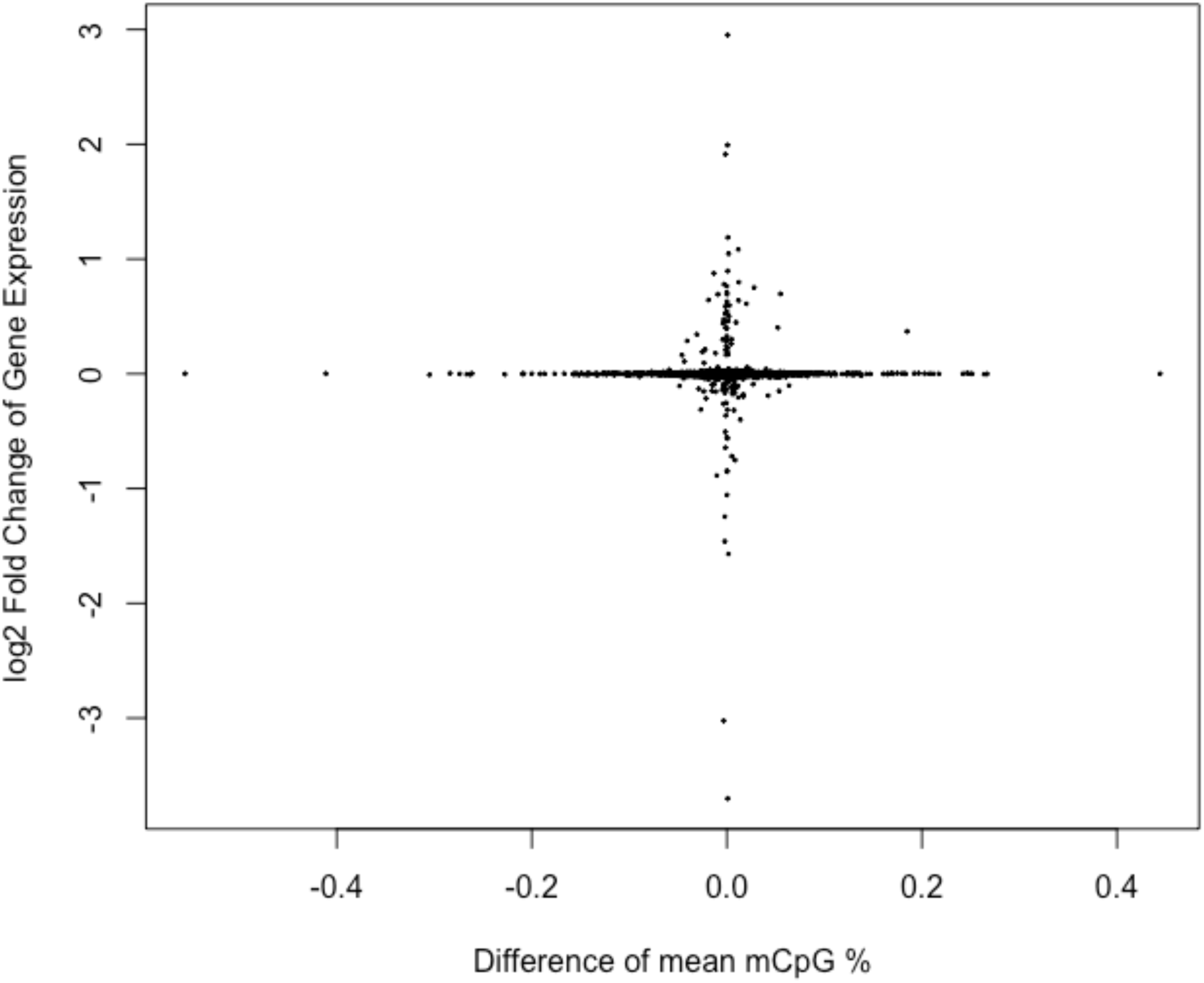
Differential methylation is not associated with differential gene expression. Scatterplot of differential expression against differential methylation. The pattern would approximate a 1:1 relationship if cytosine methylation directly influenced gene expression. Each circle represents an individual gene.

## Discussion

DNA methylation and its machinery is present within many insect species but there is variation among species for both genomic level and copy number of the genes influencing DNA methylation, *Dnmt1* and *Dnmt3* (Bewick et al. 2017). While DNA methylation itself does not have a universal function among insects, one of the genes, *Dnmt1*, consistently perturbs gametogenesis when its expression is reduced (Zwier et al., 2012; Zhang et al., 2015; Schulz et al., 2018; Bewick et al., 2019; Li et al. 2019; Amukamara et al., 2020; Gegner et al., 2020; Ventós-Alfonso et al., 2020; Takada et al., 2021; Washington et al., 2021; Arsala et al., 2022; Xu et al. 2022; Cunningham et al., 2023; Ivasky et al., 2023; Shelby et al. 2023). Here, we investigated the genetic underpinning of the severe reproductive deficits after *Dnmt1* expression was reduced using RNAi fed to adult female *B. tabaci*. This treatment, at the reduced expression level observed here, causes females to produce both fewer and less viable eggs (Shelby et al., 2023). We used RNA-seq of whole-body samples to specifically test an *a priori* hypothesis that this treatment would impact meiotic pathways preferentially, the developmental point at which many gametes appear to arrest (Amukamara et al., 202; Washington et al., 2021; Cunningham et al., 2023). We also explored this dataset to examine if other genetic pathways associated with germ cell maturation were impacted. Relatively few genes overall were differentially expressed. This work could have detected strong effects, we believe the lack of pervasive differential gene expression given the severe reproductive deficit phenotype is likely due to using whole bodies. While our hypothesis of meiosis-specific pathways was not supported, we did find differential expression of *Wnt*-like genes. From this, we think that *Wnt* and its pathways become an interesting candidate pathway for additional study.

We also performed a baseline characterization of genomic cytosine methylation of *B. tabaci* and the impact of *Dnmt1* knockdown on the level of methylation. Cytosine methylation was found at comparable level to other hemipteran, within a CpG context, and was enriched within genic regions across the genome. Cytosine methylation differences between genes were positively associated with gene expression level. However, differences of cytosine methylation level due to treatment within the same gene were not associated with gene expression differences across all genes. No statistically significant differences in gene body methylation were detected. This is likely attributable to overall few cellular divisions happening at the whole-body level where *Dnmt1*’s reduced expression is likely to manifest its effect. Moving forward, there is a need to take a tissue specific approach to understanding the genetics of this phenotype. Our analysis also provides a new candidate pathway for the study of reproductive deficits of female insects and provide a resource for studies of genomic methylation of *B. tabaci*.

Despite the dramatic reproductive phenotype, we did not find large numbers of differentially expressed genes, a lack also observed with other studies (Bewick et al., 2019; Cunningham et al., 2023). This suggests that *Dnmt1* knockdown is affecting specific genes or gene networks, rather than producing the reproductive deficit by instigating a global transcriptional response. Our *a priori* hypothesis that meiotic specific pathways would be more affected compared with mitosis or general cell cycling pathways did not receive strong support overall. None of our meiosis candidate genes were differentially expressed compared with one or two in the other categories – general cell proliferation, cell cycle, and mitosis. Genes that were annotated with the GO term of meiosis were not enriched among the genes ranked highest by expression differences, regardless of whether they differentially expressed or not. Among only the differentially expressed genes themselves (96 genes), one meiosis GO term was overrepresented. However, the list also contained several mitosis specific and development specific GO terms. We believe this general overrepresentation of development genes is an overall reflection of assaying a perturbed reproductive tissue and not specific of any pathway being preferentially affected. Thus, we did not find support for *Dnmt1* being directly involved in the control of meiosis with this study.

However, we found intriguing evidence that a conserved genetic pathway, the *Wnt* pathway, might be affected by *Dnmt1* knockdown. *Wnt*’s of insects are usually viewed as general development genes; involved with cell proliferation, migration, patterning, and identity (Murat et al., 2010; Waghmare and Page-McCaw, 2018). In human colon cancer cells, *Dnmt1* influences the *Wnt* pathway via its members *β-catenin* and *Snail1* through a DNA methylation-independent mechanism (Espada et al., 2011). While mammalian DNA methylation operates differently than it does for insects, these results generate specific, testable hypotheses about how the expression of *Dnmt1* is required for proper regulation of gametogenesis. The regulation of the *Wnt* pathway by *Dnmt1* being methylation independent also aligns with much of the evidence produced by insect systems (Bewick et al., 2019; Amukamara et al., 2020; Ventós-Alfonso et al., 2020; Washington et al., 2021; Arsala et al., 2022; Cunningham et al., 2023; Ivasky et al., 2023; Shelby et al., 2023). This pathway has also been shown to be involved in entry into meiosis of mice (Chassot et al., 2011, Le Rolle et al., 2021). Our future experiments will explore the interaction between the *Wnt/β-catenin* and *Dnmt1* in *B. tabaci*.

One possible explanation for detecting few differential expressed genes is that we sampled not only the tissue in which *Dnmt1* knockdown has its observed effect, the ovaries, but whole bodies. However, *Dnmt1* expression was reduced, and the magnitude of reduction observed here has been shown to reduce the presence its protein, DNMT1, within the nucleus of ovarian cells (Shelby et al., 2023). The observed amount of differentially expressed genes is reasonable from whole bodies of this species when the experimental effects are regional or tissue specific (Tadmor et al., 2022). A second possible explanation is that we did not detect high levels of differentially expressed genes because we did not appreciably reduce DNA cytosine methylation. The reduction in DNA methylation with loss of *Dnmt1* requires cell division and DNA synthesis. High turnover tissues exist within adult insects, such as digestive and reproductive tissues. However, they would be the minority of the cell pollution sampled here. Thus, our bisulfite sequencing data might not detect differential DNA methylation within the small proportion of cells that divided between treatment and sampling (24 hours). However, given the consistent result that changes of cytosine methylation pattern are not associated with differential expression in *O. fasciatus* gonads (Bewick et al., 2019, Cunningham et al., 2023), our data support the hypothesis that *Dnmt1* knockdown is affecting targeted genetic pathways in the gonads to specifically affect gametogenesis. This adds to the evidence that *Dnmt1* or lack thereof is acting through a cytosine methylation independent pathway. The gametes/eggs that were affected were post-meiotic and such cells would have similar cytosine methylation patterns to control samples; however, they lacked nuclear DNMT1 (Shelby et al., 2023). Recently, an ancestrally duplicated domain of DNMT3 of honey bees showed binding to histone residues, a previously uncharacterized interaction within insects (Kucharski et al., 2023). Reanalysis of the alternative splicing of this gene during development also showed many isoforms produced with the duplicated domain and without the catalytic methyltransferase domain (Kucharski et al., 2023). This supports the general idea that these *Dnmt* gene families are more diverse than previously appreciated and that they have the capacity to act via different pathways. This would suggest that *Dnmt1* knockdown effects are not likely being mediated through difference of cytosine methylation.

Our final aim of the study was to characterize the cytosine methylation of *B. tabaci*. This was both a general characterization and how *Dnmt1* knockdown affected the patterns. Cytosine methylation was found within a CpG context and enriched within genic regions. Genes have a bimodal distribution whereby gene are either highly methylated or not and methylation level is associated with gene expression level between loci. There was a positive association between methylation level and absolute level of gene expression and a negative association with gene expression variability. All of these patterns are common among insects and suggest that cytosine methylation is comparable among hempiterans (Bewick et al., 2017; Provataris et al., 2018). There was little cytosine methylation found across transposons, which suggests that this nucleotide modification is not serving as a defense against those elements (Bewick et al., 2019; Gladstad et al., 2014). The RNAi *Dnmt1* treatment produced no statistically significant differentially methylated genes. We believe the same explanation for few differentially expressed genes applies to this result as well – whole body samples were not sensitive enough to detect changes of gene methylation within a minority of cells within our samples. Although methylation level between treatments were not statistically significantly different, some loci did show differences; these differences were not associated with gene expression levels. This also makes *B. tabaci* comparable to other insects where an association between difference of gene methylation and expression is rare (Moradin and Brendel, 2021; Dixon and Matz, 2022; Duncan et al., 2022; Malezka and Kucharski, 2022). Overall, *B. tabaci* shows many of the conserved patterns of cytosine methylation of insects but a more tissue specific approach is warranted before making definitive conclusions.

Pest management of *B. tabaci* is one of the primary reasons for any work with this species. Our results here are add to the understanding of its reproduction and will inform experimental design moving forward. Clearly, *Dnmt1* knockdown produces severe reproductive deficit for female *B. tabaci* (Shelby et al., 2023). We also now know that *B. tabaci* has cytosine methylation patterns and association like other hemipteran and insects meaning we can use those species to help inform hypotheses, predictions, and conclusions. The work here is an initial attempt to fill in the mechanistic understanding of how *Dnmt1* might work. This current work also suggests that we need to focus on disrupted tissues themselves, which should facilitate a more complete understanding of the pathways and mechanisms by which *Dnmt1* influences insect reproduction.

### Experimental Procedures

To understand the functional role that *Dnmt1* plays during gamete formation of female *B. tabaci*, we used RNAi to knockdown *Dnmt1* gene expression replicating the protocol of Shelby and colleagues (2023). After the treatment, we tested two sets of *a priori* predictions after gene expression quantification using RNA-seq; one using candidate genes and the other using Gene Ontology terms. The same RNA-seq data were also used to perform exploratory analyses to assess how *Dnmt1* knockdown perturbed the transcriptome of females. We also assessed cytosine methylation of both control and RNAi-treated females. We first characterized the level *B. tabaci* has genomic methylation, in what genomic context it is found, how gene methylation related to gene expression globally. We then assessed if *Dnmt1* expression reduction caused differential gene methylation and if differential gene methylation was associated with differential gene expression.

### Animal colony and husbandry

This experiment used the same laboratory population and husbandry protocols as Shelby and colleagues (2023). Briefly, these *B. tabaci* colonies were started from population collected from a cotton (*Gossypium hirsutum*) field site in Tift County, Georgia, USA in 2018 (McKenzie et al., 2020) belonging to the MEAM1 *B. tabaci*. The whiteflies were maintained on collard (*Brassica oleracea*) plants using individual pots and grown to a minimum height of 15 cm. Both whiteflies and collards were maintained at 26°C with a 14:10 hour light:dark photoperiod in 51 cm × 28 cm × 28 cm aluminum insect cages (Bioquip Products, Rancho Dominguez, CA, USA) within a Percival Incubator (Percival Scientific, Perry, Iowa, USA) during the entire experiment.

### RNA interference (RNAi) synthesis, administration, and quality control

We created RNAi constructs targeting *Dnmt1* following Shelby and colleagues (20123). Briefly, RNAi constructs targeting *Dnmt1* were prepared via PCR using 500 ng of RNA from unmated females. The construct was targeted towards the Replication Fork Domain (RFD) motif the *Dnmt1* gene. Afterwards, double stranded RNA was synthesized with an Ambion MEGAscript kit (ThermoFisher Scientific, Waltham, MA, USA) following the manufacturer’s protocol. This reaction was purified with a phenol:chloroform:IAA extraction followed by a sodium acetate precipitation. We measured concentration using Invitrogen’s Qubit system with the ssRNA kit following the manufacturer’s protocol. We used *eGFP* as an exogenous control construct.

Newly-eclosed females were removed from colonies within 24 hours post-eclosion to drastically reduce the opportunity to mate. Thereafter, adult female at 3-6 days post-eclosion were fed 200 µl of RNAi solution of either *Dnmt1* or *eGFP* each at a concentration of 0.05 µg/mL within a 10% w/v sucrose solution contained within a cap of a 1.7 mL microcentrifuge tube using an artificial feeding apparatus made from microcentrifuge tube caps. Females were haphazardly allocated to treatment group. The RNAi solution was colored with 5 µl of green food coloring. Parafilm was placed over the cap to provide a membrane for feeding. 75–100 female whiteflies were placed into each prepared acetate tube, which was then wrapped with black construction paper to promote movement to the top of the tube. Females were allowed to feed for 24 hours, and consumption was confirmed by observing a green abdomen before collection. *eGFP* control and *Dnmt1* females were collected after this feeding period and immediately flash frozen with liquid nitrogen and stored at − 80°C until nucleotide extraction. Twenty-five females were pooled into each biological replicate.

This treatment does not impact lifespan between control and *Dnmt1* treated females or gross morphological structures (Shelby et al., 2023). This treatment statistically significantly reduces the expression of *Dnmt1* of treated females (Shelby et al., 2023).

### RNA & DNA extraction

Total RNA and genomic DNA was extracted using a Qiagen Allprep DNA/RNA Mini Kit (Qiagen, Venlo, The Netherlands) following the manufacturer’s protocol. Homogenization of the females was performed using a Bead Bug (Benchmark Scientific, Sayreville, New Jersey, USA) at 4000 rpm for 60 seconds. Quantification of DNA and RNA was done with a Qubit fluorometer using the DNA HS and RNA BR kits, respectively, following the manufacturers protocol.

### RNA-seq High-throughput library preparation and sequencing

Twelve biological replicates each of *eGFP* and *Dnmt1* were used for the RNA-seq experiment. A sample of 150 ng of extracted total RNA of *B. tabaci* was used to construct poly-A selected, non-stranded Illumina compatible libraries by Novogene Corporation (Sacramento, CA, USA). We targeted 20M 2 x 150 bp read pairs per biological replicate using an Illumina NovaSeq by Novogene.

### RNA-seq quality control and mapping

Reads were initially assessed for quality with fastQC (v0.11.9; default settings; Andrews, 2010). Reads had adapters trimmed with cutadapt (v2.8; --trim-n -O 3 -u -2 -U -2 -q 10,10 -m 30; Martin, 2011) using the TruSeq adapter sequences. Afterward, reads were reassessed with fastQC with default settings. Overlapping reads were merged with FLASh (v 1.2.11; default settings; Magoc and Salzberg, 2011). As a final QC step, reads were mapped to rRNA genes and were removed with SortMeRNA (v4.3.3; s rRNA: NC_006279, l rRNA: NC_006279.1; Kopylova et al. 2012). We used the NCBI RefSeq assembly (ASM185493v1; Chen et al., 2016) and annotation (GCF_001854935.1). HISAT2 (v2.2.1; no soft-clipping; Kim et al. 2019) was used to map reads to the genome. Mappings were converted to read counts by StringTie (v2.1.7; Pertea et al. 2015) following the manual instructions for export to DESeq2.

### Functional annotation

We added functional annotation to the *B. tabaci* proteome/transcriptome using eggNOG-mapper webserver using the Auto taxonomic scope setting (http://eggnog-mapper.embl.de; v 2.19; Cantalapiedra et al. 2021). This annotated 16,813 of 22,737 protein/transcript models (74.1%) with Gene Ontology terms, which was the database to perform our GO term enrichment analysis and GO term overrepresentation analysis (Supplementary File 1).

### *a priori* candidate gene screen

We tested our hypothesis that *Dnmt1* knockdown would preferentially influence meiotic, not mitotic, or general cell cycle, pathways by assessing a series of candidate genes associated with each pathway. These genes were chosen as canonical members based on literature searches (Cunningham et al., 2023). These include cell fate and proliferation genes that we did not expect to be differentially expressed (members of the *Wnt*, *Frizzled*, and *Pumillio* families), cell cycle control genes (some of which we did expect to be differentially expressed (*Vasa*) and some of which we did not expect to be differentially expressed (*Cyclin B3*, *Cyclin D2*, *Cyclin D3*, *Cell Division Cycle 20*, *Cell Division Cycle 25* family members)), maintenance of chromosome genes that we did expect to be differentially expressed (*Structural Maintenance of Chromosome 3* family members), meiotic transition gene that we did expect to be differentially expressed (*Boule, Decapentaplegic*, and *Glass boat bottom* families), and meiotic recombination genes that we expected to be differentially expressed (*SPO11 Initiator of Meiotic Double Stranded Breaks*, *MutS Homolog 5*, *Meiotic Nuclear Divisions 1*, and *Homologous-Pairing Protein 2* families). We did not predict specific directional changes, except decreased expression of *Vasa* and *Boule* within the *Dnmt1* knockdowns. We extracted raw P values for differential gene expression from DESeq2 and compared them with Adaptive Two-Step B-H Thresholding test, which is less conservative than standard Benjamini-Hockberg correction (Benjamini et al., 2006).

### *a priori* Gene Ontology (GO) term enrichment analysis

We also tested our hypothesis using a very broad test of pathway enrichment. We selected five high-level GO terms representing the genetic pathways that we predicted would or would not be after *Dnmt1* knockdown. These were GO:0007049 Cell Cycle, GO:0000278 Mitotic Cell Cycle, GO:0051321 Meiotic Cell Cycle, GO:0007276 Gametogenesis, and GO:0007283 Spermatogenesis. With our previous observed phenotypes, we expect Meiotic Cell Cycle, Gametogenesis, and Spermatogenesis to be enriched. We expected Cell Cycle and Mitotic Cell Cycle to show no enrichment. We included GO:0070265 Necrotic Cell Death and GO:0006915 Apoptotic Process to assess if our samples were undergoing these processes. We used the S values from DESeq2 to rank order genes. We used fgsea R package (Korotkevich et al. 2021) with default parameters to test if genes annotated with these GO terms were enriched at the head of these lists. The same R package was used to visualize the results.

### Differential gene expression

Read counts were imported into R using tximport (Bioconductor v1.20.0; Soneson et al. 2015) following the manual’s instructions. We used R (v4.1.0; R Core Team, 2021) within an Rstudio IDE (build 492; Rstudio Team, 2022) for the analysis.

DESeq2 (Bioconductor v1.32.0; default settings; Love et al. 2014) was used for the differential gene expression analyze following the manuals suggestions for exploratory data analysis and sample/analysis quality control. *eGFP* was set and used as the comparison group. We checked the model matrix specification to ensure correct specification (i.e., that program was contrasting the samples in the correct way). After importing and initial analysis, samples were plotted with a PCA to visually check for outliers according to the manual’s recommendation. After inspection, we removed one *Dnmt1* sample that was both an intra-group PCA outlier and having very high expression of *Dnmt1* that suggested the RNAi had not worked properly. This left 12 biological replicates of *eGFP* and 11 of *Dnmt1* for the final differential gene expression analysis.

After removal of the outliers the analysis was repeated using the same settings of DESeq2 and the results were again quality controlled. We used the default dispersion estimator and shrinkage method, apeglm (Zhu et al. 2019). We used s-values to estimate statistical significance after false discovery rate correction at the level of 0.05 (Stephens 2016). Results were visualized with the fviz_pca_ind function of the factoextra R package (Kassambara and Mundt 2020).

### Gene Ontology (GO) term overrepresentation analysis

We found overrepresented GO terms among the differential expressed gene to provide us with biological processes that might be perturbed using the topGO package of R (v2.48.0; Alexa et al. 2006). Often termed an enrichment analysis (Huang et al. 2009), the analysis specifically tests for overrepresentation of a GO term from a cohort of provided genes (usually those that are differentially expressed) compared with the expectation of the same number of random picks from all expressed genes. We performed these GO term tests using Fisher’s exact test with the weighted algorithm.

### Whole-genome bisulfite sequencing (WGBS) library preparation and sequencing

Five biological replicates each of *eGFP* and *Dnmt1* were used for the differential methylation analysis. A sample of 150 ng of genomic DNA was used to construct bisulfite treated genomic DNA Illumina compatible libraries by the Novogene Corporation (Sacramento, CA, USA). Lambda phage gDNA, which is known to be completely unmethylated, was added to these libraries to provide a negative control and estimate conversion efficiency. We targeted 30X coverage of the genome using 2 x 150 bp pair-end reads. Sequencing was performed on an Illumina NovaSeq.

### WGBS quality control, mapping, and cytosine methylation analysis

As described above, reads were quality assessed with fastQC, trimmed with cutadapt using the TruSeq adapter sequences, and reassessed with fastQC with default settings. Reads were mapped to the NCBI RefSeq assembly (ASM185493v1) & annotation (GCF_001854935.1) using HISAT-3N (v2.2.1; C to T base change, uniquely mapped non-clonal reads; Zhang et al., 2021; Supplementary File 2). We then used HISAT-3N to generate a per nucleotide results table for all cytosines and a separate one for cytosines within a CpG context, which is were almost all of cytosine methylation is found for insects (Bewick et al., 2017). Using the HISAT-3N results tables per sample, we first filtered out cytosines that had fewer than five mapped reads. We then used Python and the numpy, scipy. stats, statsmodel.stats, and pandas libraries to generate per nucleotide result tables of percent reads that indicated methylation, total mapped reads, and one-tailed binomial tests with FDR-corrected P values for each sample (P < 0.016, which was the average non-conversion rate of reads mapped to the lambda phage controls). Methylation status of each cytosine was collected into a per gene results table for all samples using bedtools (Quinlan and Hall, 2010) and pandas. We also generated a results table that contained only CpGs that were symmetrically mapped above our threshold. To be able to characterize cytosine methylation over repeats and transposons, we identified transposable elements using RepeatMasker (v4.1.2; Smit et al., 2013) which was consistent with the invertebrate repeat library from Repbase (Jurka, 1998) following Bewick and colleagues (2019). Cytosine methylation over these genomic regions was collated with bedtools.

### Characterization of cytosine methylation

We characterized the genomic context of cytosine methylation of the *B. tabaci* in several ways. We first characterized the GC content of the genome using bedtools. We then calculated the percent of cytosines that were methylated for several genomic regions: the genome as a whole, intragenic regions, genes, exons, and introns. Lastly, we characterized percent of cytosines methylated across genes and transposons with bedtools and plotted with deepTools2 (v3.5.1; Ramirez et al., 2016).

### Differential gene methylation and analysis

After understanding the genomic context of cytosine methylation, we focused on understanding its association with gene expression. First, we expected to see an association between gene methylation and gene expression such that highly methylated genes were also the highest expressed genes (Duncan et al., 2022). To that end, we divided genes into deciles and plotted those values against the mean standardized expression of the gene within each decile using R. We also plotted how differences of gene methylation were associated with gene expression standard deviations.

To understand the effect of *Dnmt1* RNAi on gene methylation, we tested for statistically significantly differences of gene methylation levels between the two treatments using the t.test function of R. Before this test, we filtered the results for genes that showed at least a 1% difference for percent of cytosines methylated (2,391/13,901 genes) so the *t* test would not return an error due to data showing no variation. We plotted the mean percent cytosines methylated for each gene between the two treatments.

Lastly, we assessed how differences of gene methylation were associated with difference of gene expression by plotting log2 Fold Changes of gene expression by difference of mean percent methylated cytosines.

## Supporting information

Updated B tabaci Gene GO Terms

Differentially Expressed Gene SI

GO Term Overrepresentation SI

## Acknowledgements

We appreciate the discussions and contributions from other members of our laboratories, especially Ahva Potticary and Bazgha Zia. The mention of a proprietary product does not constitute an endorsement or a recommendation for its use by USDA or University of Georgia.

## Author Contribution Statement

CBC, PJM, AJM, EAS, and AS designed the study. EAS proposed examining the *Wnt* genetic pathway. EAS and ECM performed the experiments. CBC analyzed the data. CBC, PJM, AJM, and EAS interpreted the data. All authors contributed to drafting the manuscript.

## Data Availability

All high-throughput data is available under NCBI BioProject # XXXX. Analysis scripts are available at github.com/cbcunningham/Bt_RNA-seq_methylC-seq.

